# FarmGTEx TWAS-server: an interactive web server for customized TWAS analysis in both human and farm animals

**DOI:** 10.1101/2023.02.03.527092

**Authors:** Zhenyang Zhang, Zitao Chen, Jinyan Teng, Shuli Liu, Qing Lin, Yahui Gao, Zhonghao Bai, The FarmGTEx Consortium, Bingjie Li, George Liu, Zhe Zhang, Yuchun Pan, Zhe Zhang, Lingzhao Fang, Qishan Wang

## Abstract

Transcriptome-wide association study (TWAS) is a powerful strategy for elucidating the molecular mechanisms behind the genetic loci of complex phenotypes. However, TWAS analysis is still daunting in many species due to the complication of the TWAS analysis pipeline, including the construction of the gene expression reference panel, gene expression prediction, and the subsequent association analysis in the large cohorts of genome-wide association study (GWAS). Farm animals are major protein sources and biomedical models for humans. To facilitate the translation of genetic findings across species, here we provide an interactive and easy-to-use multi-species TWAS web server for the entire community, called the FarmGTEx TWAS-server (http://twas.farmgtex.org), which is based on the GTEx and FarmGTEx projects. It includes gene expression data from 49, 34, and 23 tissues in 838 humans, 5,457 pigs, and 4,889 cattle, representing 38,180, 21,037, and 17,942 distinct eGenes in prediction models for humans, pigs, and cattle, respectively. It allows users to conduct gene expression prediction for any individuals with genotypes, GWAS summary statistics imputation, customized TWAS, and popular downstream functional annotation. It also provides 479,203, 1,208, and 657 tissue-gene-trait association trios for the research community, representing 1,129 human traits, 41 cattle traits, and 11 pig traits. In summary, the FarmGTEx TWAS-server is a one-stop solution for performing TWAS analysis for researchers without programming skills in both human and farm animal research communities. It will be maintained and updated timely within the FarmGTEx project to facilitate gene mapping and phenotype prediction within and across species.

## INTRODUCTION

Genome-wide association studies (GWAS) have discovered numerous genetic variants associated with complex diseases and traits in both human and livestock populations (1-4). However, most of these variants are in high linkage disequilibrium (LD) and reside in noncoding regions, which makes it extremely challenging to interpret their underlying molecular mechanisms. Integration of multi-omics data has been proven to be efficient in understanding the mechanisms of action of noncoding variants behind complex phenotypes. Among those methods, transcriptome-wide association study (TWAS) is a popular one (5). In brief, TWAS first derives the gene expression prediction models by using a regression model or non-parametric approaches from a reference panel with both genotype and gene expression. With these prediction models, gene expression levels of individuals in GWAS populations can be predicted based on their genotype data. And then we can associate the predicted expression levels (the genetically controlled proportion) of each gene with the phenotypes of interest (5). To date, various TWAS software packages have been developed, e.g., PrediXcan/S-PrediXcan (5), TWAS FUSION (6), UTMOST (7), MR-JTI (8), TIGAR (9), and PUMICE+ (10).

In human genetics, projects like Genotype-Tissue Expression (GTEx) (11) provided a valuable gene expression reference panel across various tissues in hundreds of individuals and paved the way to systemically characterize the regulatory effects on complex traits and diseases *via* TWAS (12-17). Several webservers for TWAS analysis and results sharing such as webTWAS (18), TWAS-hub (19) and TWAS atlas (20) are available in human. However, TWAS studies in livestock lag far behind humans. The Farm animal Genotype-Tissue Expression (FarmGTEx, https://www.farmgtex.org/) project has been established to provide the transcriptome reference panel across a wide range of tissues in farm animal species, including cattle (21), pig (22), and other species in the future. Although the genotype and gene expression data are available, the complicated TWAS analysis is still challenging and time-consuming for most of the researchers who do not have a solid background in bioinformatics and statistical genetics. In addition, translating genetic findings across species is also important in the field of evolution, biology, and genetics. For instance, previous studies demonstrated the conserved functional impacts of orthologous variants on gene expression and complex traits between livestock and humans (23-28). Liang et al. (29) also proposed polygenic transcriptomic risk scores (PTRS), which paved the way to translate polygenic signals across human ancestry groups. Moreover, livestock has been proposed as a desirable model for human biology and medicine studies. For example, the pig shows more similar body size, organ size, physiology, and anatomy to humans (30), which makes it a suitable biological model used for drug design and organ xenotransplantation in human medical research(31,32). Therefore, combining genetics studies in humans and farm animals will become crucial and worthwhile for understanding the molecular and evolutionary basis of complex phenotypes across species.

In this study, to facilitate the TWAS analysis and the translation of genetic findings across species, we develope the FarmGTEx TWAS-server, which is the first user-friendly web server that allows users to conduct the TWAS analysis across multiple species including human, pig, and cattle. We implement three popular and classical TWAS software packages: S-PrediXcan (5), TWAS-FUSION (6), and UTMOST (7). By uploading the GWAS summary statistics, users could perform the TWAS analysis conveniently. We also provide functions including liftOver, GWAS summary statistics imputation, gene set enrichment analysis (GSEA), and result visualization. Incidentally, we provided summary statistics of TWAS from many complex traits in humans, cattle, and pigs. The FarmGTEx TWAS-server is an open-access resource that is freely available at http://twas.farmgtex.org, and it will be updated timely and include more species as the FarmGTEx project is expanding.

## MATERIAL AND METHODS

### Gene expression data collection and normalization

The expression (Transcripts per Million, TPM) of 26,908 and 27,537 genes from 34 and 23 tissues in pigs and cattle were obtained from the FarmGTEx project (21,22), respectively. Details of these samples are summarized in Supplementary Table 1 and Supplementary Table 2. For each of the tissues in pigs and cattle, genes with TPM < 0.1 and raw read counts < 6 in more than 20% of samples were excluded. Finally, a total of 5,457 and 4,889 samples were analyzed in pigs and cattle, respectively. Gene expression values were sample-wise corrected using the trimmed mean of M values (TMM) (33), followed by the inverse normal transformation of TMM. More details have been reported in (21,22). The TPM of 55,878 genes from 54 human tissues were downloaded from the Genotype-Tissue Expression (GTEx) project (https://www.gtexportal.org/) (11), among which five tissues were excluded due to the small sample size, including bladder (n=21), cervix_ectocervix (n=9), cervix_endocervix (n=10), fallopian_tube (n=9), and kidney_medulla (n=4). The sample size of all the human tissues used in the TWAS sever is summarized in Supplementary Table 3.

### Gene expression prediction model training

To build gene expression prediction models based on the transcriptome reference panels, we used the Elastic Net model in S-PrediXcan(5), Top1, BLUP, and BSLMM models in FUSION (6), and CTIMP in UTMOST (7). For humans, the prediction models were downloaded from https://zenodo.org/record/3519321/ (Elastic Net model), http://gusevlab.org/projects/fusion/ (TWAS-FUSION), and https://zenodo.org/record/3842289 (UTMOST). For pigs and cattle, the Elastic Net models are available at https://www.farmgtex.org/. We further built the prediction models by using FUSION (Top1, BLUP, and BSLMM) and UTMOST (CTIMP) in pigs and cattle following the pipeline as did in humans (6,7), and the detailed parameters were referred to (21) and (22). In brief, to account for hidden batch effects of transcriptome-wide variation in gene expression within each tissue, ten PEER factors were estimated by the Probabilistic Estimation of Expression Residuals (PEER v1.3) (34) method based on the gene expression matrix. To account for the population structure, genotype PCs were estimated using PLINK v1.9 (35) based on the genotype data. The number of genotype PCs was included according to sample size: five PCs for tissues with < 200 samples, and ten PCs for tissues with ≥ 200 samples. The *cis*-window of a gene was defined as 1Mb up- and down-stream of its TSS. The prediction models of FUSION were then calculated using the Rscript FUSION.compute_weights.R --bfile $OUT --tmp $OUT.tmp --out $FINAL_OUT --verbose 0 --save_hsq --PATH_gcta $GCTA -- PATH_gemma $GEMMA --PATH_plink $PLINK2 --models top1,blup,bslmm –covar $TISSUE.covariates4Fusion.txt --crossval 5. For CTIMP models, we followed the command from https://github.com/yiminghu/CTIMP.

### GWAS data collection and quality control

To demonstrate the usefulness of this TWAS-server, we collected GWAS summary statistics of 1,129 human traits from GWAS Catalog (36), webTWAS (18), and Neale Lab UKBB v3 (http://www.nealelab.is/uk-biobank), 7 pig traits (37) and 41 cattle traits (38). Besides, we also included GWAS results of four pig traits using our newly generated genotypes of 2,778 Duroc pigs. Briefly, we genotyped these Duroc pigs with a Neogen GGP 50 K Porcine v1 Genotyping BeadChip (n = 974) or low coverage whole genome sequencing (depth = 1X, n = 1,804). We then used beagle v5.4 (39) to impute missing genotypes with the current version of Pig Genomics Reference Panel (PGRP v1) from the PigGTEx, which contained whole-genome sequence data of 1,602 pigs from over 100 breeds worldwide (22). We then performed the GWAS for four traits using GEMMA (40), including birth weight (BW), corrected days to 115 kg (DAY115), back-fat thickness correct for 115 kg (BFT115), and loin muscle area corrected for 115 kg (LMA115).

We only considered the GWAS summary statistics with full information including dbSNP ID or variant coordinate, effect/non-effect allele, *P*-value, beta coefficient, and z-score. To ensure the format of GWAS data acceptable by TWAS software, we performed the following quality control. 1) In the human dataset, for variants only with variant coordinates, we retrieved their rsID from dbSNP build 151. While in animal datasets, we used the variant coordinates for TWAS based on Sscrofa11.1/susScr11 or ARS-UCD1.2/bosTau9 for pigs and cattle, respectively. If the GWAS summary statistics were based on different genome assemblies, we performed the liftOver analysis by PyLiftover v0.4 (https://pypi.org/project/pyliftover/). 2) We removed the GWAS summary statistics that the non-effect allele or effect allele wasn’t clearly determined. 3) We excluded the GWAS summary statistics without *P*-value and beta coefficient. 4) After performing TWAS, we discarded the results with less than ten genes being tested.

### Imputation module for GWAS summary statistics

To enhance the power of TWAS, we constructed the GWAS summary statistics imputation module. The genome reference panels were obtained from 1000 Genomes (41), CattleGTEx (21), and PigGTEx (22) for human, cattle, and pig, respectively. In the “GWAS imputation” module, we provided the whole genome sequence panel including 27,731,499 (n = 500), 3,824,445 (n = 7,394, the variants were called from RNA-seq data), 42,523,218 (n = 1,602) variants for humans, cattle, and pigs, respectively. To reduce the computational burden, we only considered SNPs identified as significant variants in the gene expression prediction model (eVariants) for the “GWAS imputation”. The number of eVariants for each prediction model is summarized in Supplementary Table 5.

We considered two software packages for GWAS summary statistics imputation: 1) The Python-based software package summary-gwas-imputation proposed by Barbeira et al. (42), and 2) C++-based DIST (Direct imputation of summary statistics for unmeasured SNPs) (43). However, DIST did not support the imputation of cattle GWAS originally, because it did not allow the chromosome number bigger than 22. The summary-gwas-imputation could be used for three species. As they use different formats of genome reference panels as input, we constructed the respective panels for them following the pipelines in https://github.com/hakyimlab/summary-gwas-imputation and https://github.com/dleelab/dist. The human reference panels were also downloaded using the links above. In addition, we used chromosome with the largest number of SNPs to evaluate the accuracy of imputation (i.e., Pearson correlation coefficient between the imputed z-score and the observed) using the five-fold cross-validation approach.

### The workflow for online TWAS analysis

To provide a comprehensive and user-friendly TWAS web server, we allow users to perform quality control, liftOver, GWAS summary imputation, TWAS analysis, and gene set enrichment analysis (GSEA) with only uploading the GWAS summary statistics. All results and publication-quality figures are downloadable. The GWAS summary statistics file uploaded should include columns of the chromosome, position, SNP name, effect allele, non-effect allele, *P*-value, and beta coefficient. For quality control, the server will check the reference assembly, SNP, and chromosome. If the reference assembly of GWAS does not match that of GTEx or FarmGTEx, the server will use PyLiftover 0.4(https://pypi.org/project/pyliftover/) to converse the genomic coordinates. The GWAS imputation has been described above. For TWAS analysis, we prepared multiple software packages for users to choose from, including two single-tissue TWAS methods (i.e., S-PrediXcan (5) and TWAS-FUSION (6)) and a multi-tissue TWAS method (UTMOST(7)). To explore the function of a list of gene, users could perform GSEA using clusterProfiler (44). When the job is finished, we will send out an email with a link to all the job processes and results. Moreover, for users who have individual-level data, we also provide the “Expression Prediction” module, which is based on the PrediXcan (45) software package. With the imputed gene expression level, users could then quantify the association of the genetically regulated levels of gene expression with phenotypes of interest.

We provided 2,268, 41, and 15 TWAS summary statistics based on S-PrediXcan (5) for humans, cattle, and pigs, respectively, and built a user-friendly interface for users to search/query the results. In humans, due to the high density of SNP in GWAS, we conducted TWAS using the GWAS summary statistics directly. Whereas, in pigs and cattle, we conducted the GWAS imputation first and then performed TWAS analysis. Significant disease/trait**-**tissue-gene associations are defined as genes with *P*-value less than a cutoff threshold set to 0.05/n, where n is the number of genes being tested in a TWAS.

### Database and TWAS web server

We built the back end of TWAS-server using the PHP-based ThinkPHP5.0 web framework (https://www.thinkphp.cn/) and developed the front end using the Layui framework (https://github.com/layui/layui) and jQuery JavaScript library (https://jquery.com/). We established the database based on MySQL software. We used Python and R to develop the computational pipelines in the TWAS-server and visualize all data and results using the ggplot2(46) in R(47).

## RESULTS

### Overview of the FarmGTEx TWAS-server

Figure 1 shows the overview of the FarmGTEx TWAS server. Based on the gene expression reference panels in the FarmGTEx (21,22) and Human GTEx (11) projects, we first trained the gene expression prediction models in single- and multi-tissue manners. In general, the TWAS-server can take GWAS summary statistics, individual genotype and phenotype as input. As a result, it will output predicted gene expression and TWAS results. It also supports to explore the newly generated TWAS results with existing ones in the FarmGTEx server.

**Figure 1.**
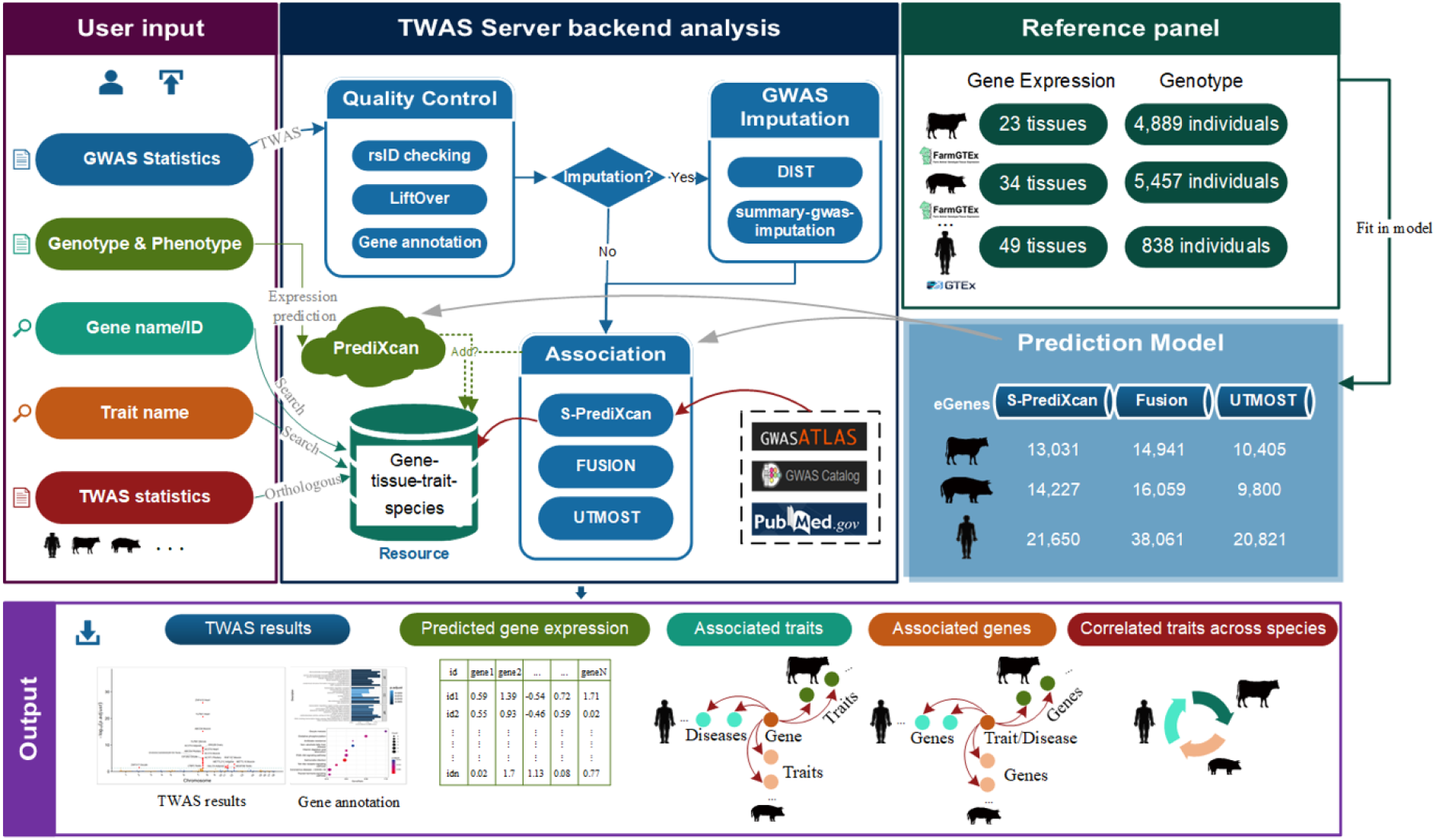
Graphical Abstract. FarmGTEx TWAS-server (http://twas.farmgtex.org) workflow. The *Reference panel* represents the gene expression data & genotype data used for the gene expression prediction model. The *Prediction Model* shows the number of eGenes used in corresponding software in all the species. The color of the boxes in the *User input* is the same as that in the corresponding result in the *Output*. The name in the line connecting the *User input* and *TWAS Server backend analysis* is the corresponding module name. And the end of the arrow is the software or dataset based on. Briefly, the TWAS-server can take GWAS summary statistics (blue), individual genotype (green), gene name/ID (cyan), trait name (orange-red), and TWAS summary statistics (brown) as input. As a result, it will output the TWAS results (blue), predicted gene expression (green), traits associated with quired genes (cyan), genes associated with quired traits (orange-red), and the correlated traits based on TWAS summary statistics (brown), respectively.

### Prediction models of gene expression

In the FarmGTEx TWAS server, we provided gene expression prediction models with S-PrediXcan (Elastic Net methods), FUSION (Top1, BLUP, BSLMM), and UTMOST (CTIMP) for each of 34, 23, and 49 tissues in pigs, cattle, and humans, respectively. The sample size, eGenes (genes with significant eQTL), and eVariants of each tissue are summarized in Supplementary Table 1-3. The number of distinct eGenes and eVariants used in S-PrediXcan, FUSION, and UTMOST is displayed in Supplementary Table 4 and Supplementary Table 5. The average of estimated *cis*-heritability of genes and prediction performance of models (the square of Pearson correlation (*R*^*2*^) between predicted and observed expression in the five-fold cross-validation) are shown in Supplementary Table 6-8. We provided a total of 38,180, 21,037, and 17,942 distinct eGenes in humans, pigs, and cattle, respectively. It represented 13,780, 13,444, and 13,442 one-to-one orthologous genes in human *vs*. pig, human *vs*. cattle, and cattle *vs*. pig, respectively. Furthermore, Supplementary Table 9-11 shows the comparison of eGenes between species in terms of *cis*-heritability. The correlation of heritability ranged from 0.0921 to 0.2041 across tissues between humans and pigs. Among these tissues, the human heart (left ventricle) (n = 432) and pig heart (n = 165) showed the highest correlation (Pearson *r* = 0.20, *P*-value = 9.35E-04) with 260 shared orthologous genes being tested. Human skeletal muscle (n = 803) and pig muscle (n = 1,322) had a heritability correlation of 0.14 (*P*-value = 9.98E-07) with 1,252 orthologous genes being tested (Supplementary Table 9). In the comparison between humans and cattle, the correlation of heritability ranged from -0.0071 to 0.0733 (Supplementary Table 10).

### GWAS imputation module

To improve the power of TWAS, particularly in farm animals where GWAS are often conducted with low-or high-density SNP array, we provided the “GWAS imputation” function for imputing the GWAS summary statistics to the GTEx sequence level (i.e., matching SNPs in the eQTL mapping reference population) according to the genotype imputation reference panel from the GTEx projects. The pig genotype imputation reference panel consists of 1,602 samples with 42,523,218 variants which were generated from the whole genome sequence. The cattle genotype imputation reference panel consists of 7,394 samples with 3,824,445 variants, which were generated from the RNA-seq data. The human genotype imputation reference panel consists of 500 European individuals with 27,731,499 variants. The GWAS imputation module allows users to perform harmonization, format standardization, missing data imputation, five-fold cross-validation, and result visualization. The ‘GWAS Imputation’ tab contains the following two steps. *Step 1* allows the user to enter the “Email”, which is used to send a result link from the server (Figure 2A). Users must select one of the species and the genome assembly version, and then the server will perform liftOver if the genome reference version is different from those of FarmGTEx or GTEx (GRCh38/hg38, Sscrofa11.1/susScr11, ARS-UCD1.2/bosTau9). Users can also select the software used for imputation, including summary-gwas-imputation (42) and DIST (43). After uploading the file in the compressed format (.gz) to the server, the header of the file will be extracted, and the user must select the name for each column in *Step 2* (Figure 2A). In addition to text results, the server provides downloadable publication-quality figures (Figure 2B). Furthermore, it will output the imputation accuracy of the GWAS summary statistics using the five-fold cross-validation based on the longest chromosome (Figure 2B).

**Figure 2.**
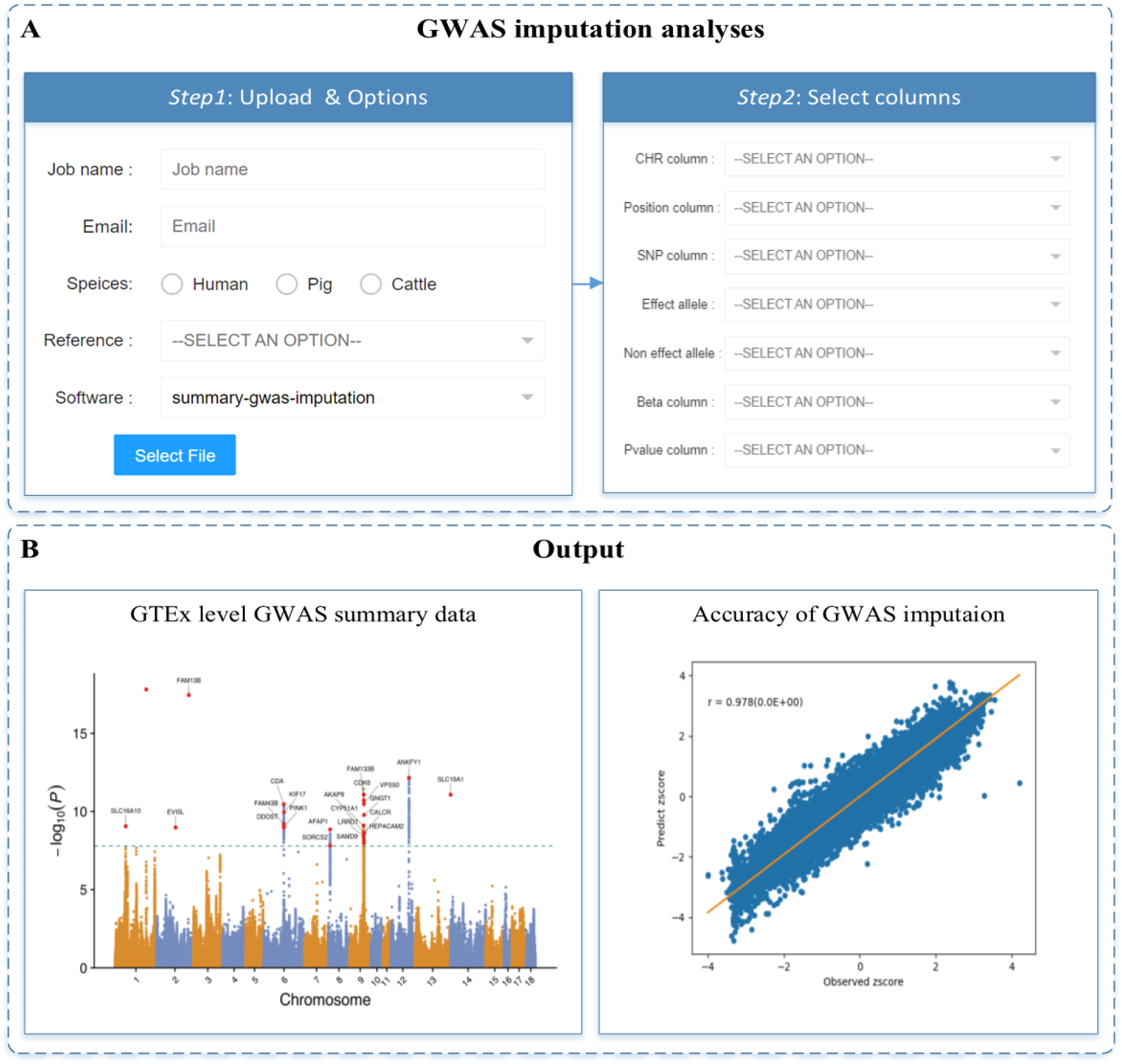
Operation flow for the ‘GWAS imputation’ module. A. *Step 1*: Upload the GWAS summary statistics and select the options. *Step 2*: Select the corresponding columns name by clicking the mouse. B. An example output for the impute GWAS summary statistics and imputation accuracy by fivefold cross-validation

### Online TWAS analysis module

The TWAS module is the major part of the FarmGTEx TWAS-server. It aims to provide a user-friendly web server for the research community to conduct the TWAS analysis easily across tissues and species based on the FarmGTEx, HumanGTEx, and other similar efforts. In the current version, it includes humans, pigs, and cattle. It will include more animals, e.g., chickens, sheep, and goats in the future, as the FarmGTEx project is working on these species. Like the GWAS imputation module, users have to upload the GWAS summary data file in *Step 1* and select the columns’ names in *Step 2* (Figure 3A). In detail, in *Step 1*, users can choose whether to do GWAS imputation in “Mode”. It will impute the genetic variants involved in the gene expression prediction models to reduce the computational time (Supplementary Table 5). Users can select different software packages to do TWAS analysis, including MetaXcan(S-PrediXcan) (5), FUSION (6), and UTMOST (7). As MetaXcan is computationally fast and has been applied in many research projects, we recommend users to try it first. To speed up the FUSION, we modified the code to allow it to run TWAS in parallel. For UTMOST, we used S-PrediXcan for the single-tissue TWAS first with the CTIMP prediction model and then performed a joint GBJ (generalized Berk-Jones) test for all the TWAS summary statistics. *Step 2* allows users to select multiple tissues to do TWAS analysis (up to 49, 34, and 23 tissues for humans, pigs, and cattle, respectively).

**Figure 3.**
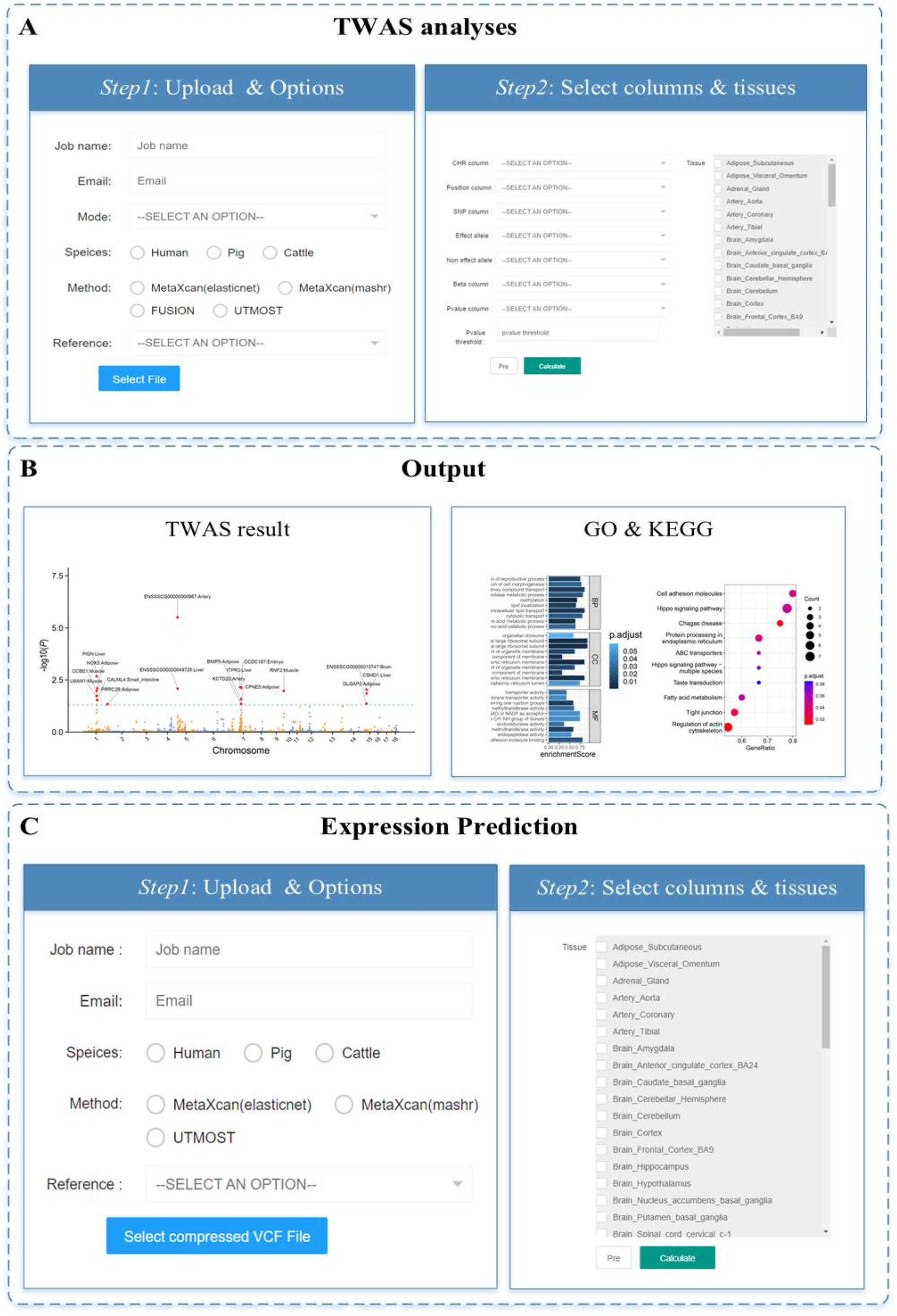
Operation flow for the ‘TWAS analyses’ module. A. *Step 1*: Upload the GWAS summary statistics and select the options. *Step 2*: select the corresponding columns name by clicking the mouse and tissues used for TWAS analysis. B. The output of the TWAS analysis. We can provide the Manhattan plots that combined all the TWAS results from multiple tissues and the visualization of the GO & KEGG enrichment analysis. C. Operation flow for the ‘Expression prediction’ module.

Users can also specify the *P-*value cutoff (default is 0.05), and the statistical significance will be defined as 0.05/n, where n is the number of genes being tested. Upon job submission, the TWAS module will perform the following six steps. (i) Quality control, (ii) LiftOver, (iii) GWAS imputation, (iv) TWAS analysis, (v) Manhattan plot illustration, (vi) GSEA for GO & KEGG enrichment analysis of genes, and (vii) result visualization. We will provide a link recording all the processes and results by *Step 3* (Supplementary Figure 1A) and send an email containing the link when the job is finished. Finally, five kinds of Manhattan plots (PDF format) can be downloaded directly, including (1) figures for GWAS (Supplementary Figure 2A), (2) figures for imputation GWAS (Supplementary Figure 2B), (3) figures for *P*-value (Supplementary Figure 2C) and z-score (Supplementary Figure 2D) of TWAS result per tissue, (4) figures for *P*-value of TWAS results from all tissues (Figure 3B), and (5) users can also zoom in a specific genomic region by the “post-Manhattan” tab (Supplementary Figure 2E). In addition to using GWAS summary statistics for TWAS analysis, the “Expression Prediction” module (Figure 3C) also allows users to predict the gene expression based on the individual-level genotype data. For doing this, users should upload the individual genotypes in VCF (compressed in gz) format, and the server will predict the gene expression level for each individual across tissues using the PrediXcan (45). We will keep users’ data on the server for a week. Afterward, we will remove the data from the server completely.

### Search module

The FarmGTEx TWAS-server curated TWAS results for 1,129 distinct human traits and diseases based on 2,268 GWAS summary data, representing 479,203 significant disease-tissue-gene trios (*P-*value < 0.05/n, n > 10). Furthermore, we added TWAS results of 41 and 7 complex traits in cattle (38) and pigs (37), respectively. Users can search these TWAS results by querying a gene or a certain disease/trait (Supplementary Figure 3). The web will provide detailed information on the quired gene and orthologous gene in other species (Supplementary Figure 3A), including the location of the gene, homology type, and ortholog confidence. It will also display the number of the disease/trait-tissue associated with the quired gene (Supplementary Figure 3B). By clicking on the digital, the corresponding disease/trait association statistics of TWAS result will show up, including gene Ensembl ID, gene symbol, trait, tissue, *P*-value, and z-score (Supplementary Figure 3C). Moreover, it will provide the gene expression of orthologous genes across tissues in all the available species (Supplementary Figure 3D). As shown in Supplementary Figure 4, the web also allows users to search for TWAS results by a particular disease/trait. It will provide detailed information about the trait, including disease/trait name, sample sizes, population, publication information, source links, and the number of associated tissue-genes detected by S-PrediXcan (Supplementary Figure 4A). By clicking on the digital, the details of the TWAS result will show up, including gene ID, gene symbol, tissue, *P*-value, z-score, and the number of associated genes in each of the tissues (Supplementary Figure 4B, C). In summary, users can explore the molecular mechanisms behind a gene or a trait based on the large-scale TWAS results, which will be a valuable resource for translating genetic findings across species.

### Cross-species mapping module

In the “Orthologous” module, users can upload TWAS summary statistics containing Ensembl ID, *P*-value, and z-score from any species such as human, pig, cattle, sheep, chicken, and mouse. The web allows users to select species and tissues to compare (Supplementary Figure 5A). Then, it will calculate the Pearson correlation coefficient based on the *P*-value and z-score of one-to-one orthologous genes between species. These results will help translate the genetic findings between species and enhance the understanding of the evolutionary basis of a particular trait/disease (Supplementary Figure 5B, C).

### Comparison to others

Currently, the FarmGTEx TWAS-server is the only web server that allows users to perform TWAS analysis across multiple species, including pigs, cattle, and humans. Compared to webTWAS (18) and TWAS-hub (19), the FarmGTEx TWAS-server is a more comprehensive server with multiple advantages, (i) more species: the webTWAS and TWAS-hub only focus on humans, whereas TWAS-server provides TWAS analysis and TWAS summary statistics across humans, cattle and pigs. In the future, we will include other farm animal species in the ongoing FarmGTEx project, such as chickens, goats, and sheep. In addition to gene expression, we will incorporate more molecular types, such as alternative splicing, promoter usage, and enhancer expression. (ii) more software packages: TWAS-hub only implements TWAS-FUSION, and webTWAS (http://www.webtwas.net/#/twas) only provides S-PrediXcan and TWAS-FUSION for online TWAS analysis. In the FarmGTEx TWAS-server, apart from the single-tissue TWAS method, we implement the multi-tissue TWAS method, UTMOST, and allow users to perform liftOver and GWAS summary statistics imputation. (iii) more functional annotations: neither TWAS-hub nor webTWAS do not allow users to perform function annotations of genes, whereas the FarmGTEx TWAS-server provides the GSEA analysis. (iv) more illustrations: apart from Manhattan plots of TWAS results, we also provide illustrations for uploaded GWAS summary statistics, imputed GWAS summary statistics (Supplementary Figure 2E).

#### Case study

Here, we presented two case studies to show how the FarmGTEx TWAS-server can help researchers to perform TWAS analysis and discover the molecular mechanisms underlying complex traits across species.

#### Case study1

We obtained the GWAS summary statistics from a previous study by Yang et al. (37), where they identified that the *ABCD4* gene was associated with total teat number (TTN). Through performing TWAS analysis using the FarmGTEx TWAS-server based on their GWAS summary data, we found that the gene expression of *ABCD4* in muscle and pituitary tissues was specifically and significantly associated with TTN (Figure 4A), suggesting the important role of muscle and pituitary in regulating TTN. In addition, due to the increased statistical power of TWAS compared to GWAS, we also found that *ABCD4* was significantly associated with left teat number (LTN) (muscle and pituitary) (Figure 4B) and right teat number (RTN) (brain, frontal cortex, muscle, blood, and small intestine) (Figure 4C). As *ABCD4* showing different *P* significance in different tissues for teat number (TN) (Figure 4), we explored the correlation between different TN traits across tissues (Figure 4D-K) and found that TTN is most correlated with LTN (*r* = 0.7-0.71), followed by RTN (*r* = 0.69-0.71) and RTN (*r* = 0.33-0.36), which is constant across tissues. *Case study1* demonstrated that the FarmGTEx TWAS-server could not only help provide regulatory mechanisms underlying GWAS loci by identifying the associated genes in relevant tissues but also increase the statistical power of association tests potentially by coming multiple signals of variants into a single gene.

**Figure 4.**
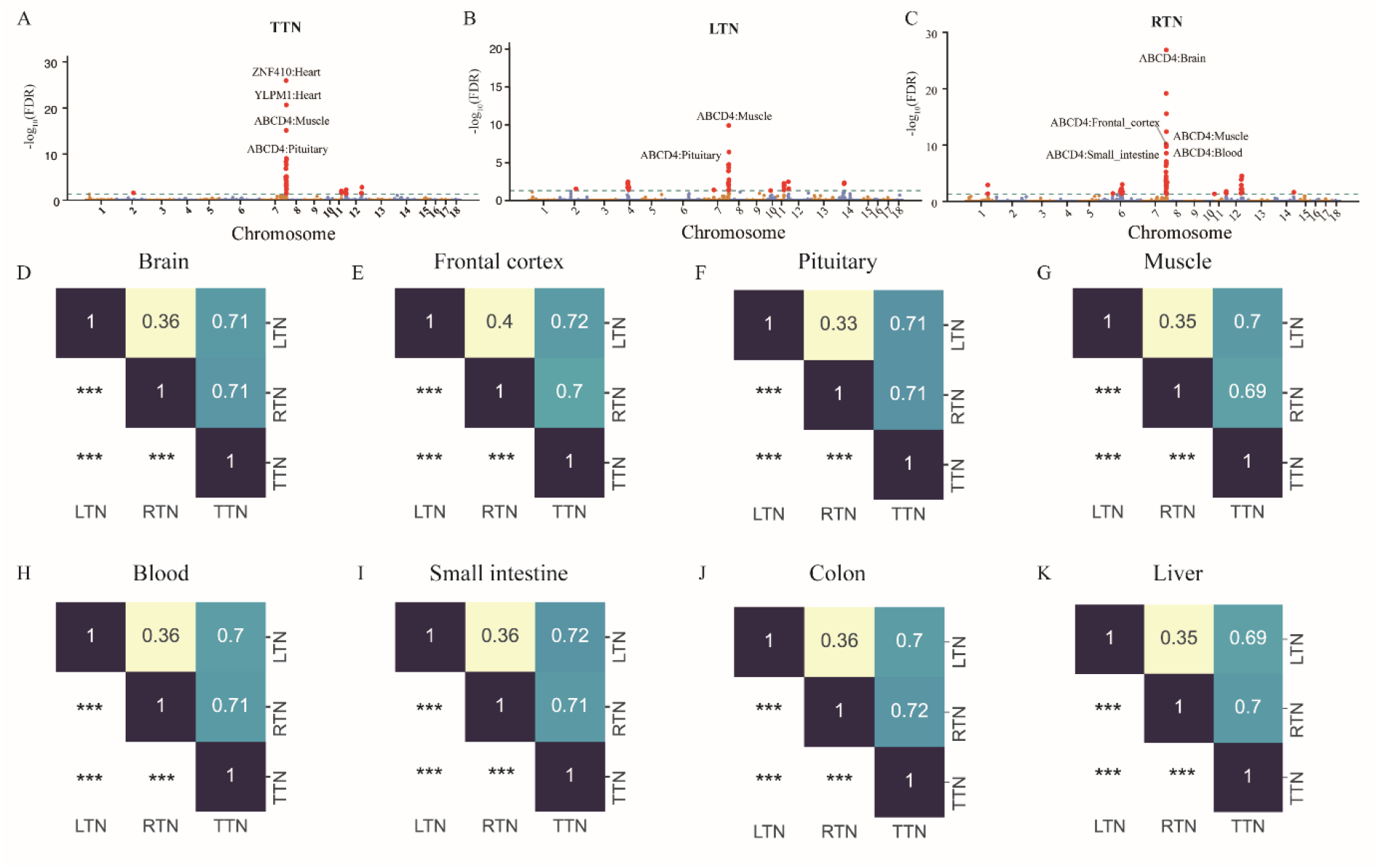
Results for *Case study 1*, TWAS results calculated from the published paper. The Manhattan plots of TWAS results for TTN(A), LTN(B), and RTN(C). FDR: false discovery rate. The upper triangle of the heatmap showed the Pearson correlation coefficient across traits in the brain(D), frontal cortex(E), pituitary(F), muscle(G), blood(H), small intestine(I), colon(J), liver(H). The lower triangle of the heatmap represents the statistical significance of the Pearson correlation. *** represents Pearson correlation *P* value < 0.001.

#### Case study2

To illustrate the usefulness of the FarmGTEx TWAS server in translating genetic findings across species, we compared the TWAS summary statistics of body conformation-related traits in humans, pigs, and cattle. It included body height (BH), body weight (BW), and body mass index (BMI) in humans obtained from Barton et al. (48), Backman et al. (49), and Sakaue et al. (50). Cattle body conformation traits includes stature, udder depth, fore udder attachment, strength, rump width, feet and legs, rump angle, body depth (38). Pig carcass traits included back fat thickness at 115 kg (BFT115), and loin muscle area at 115 kg (LMA115). They are all complex traits, and it is still challenging to elucidate their underlying mechanisms. For TWAS results of BMI, we found that *P*-value in ‘Adipose_Visceral_Omentum’ and ‘Adipose_Subcutaneous’ tissues were highly correlated with those in liver, stomach and all the intestinal tissues (i.e, esophagus muscularis, esophagus gastroesophageal junction, colon transverse, colon sigmoid, small intestine terminal ileum) (Supplementary Figure 6). Figure 5A displayed *P*-values of orthologous genes that were significantly associated between human body conformation traits and pig carcass traits in the digestive system (esophagus muscularis, esophagus mucosa, esophagus gastroesophageal junction, colon transverse, colon sigmoid, small intestine terminal ileum, and liver of human. And small intestine, large intestine, duodenum, colon, ileum, jejunum, and liver of pig). Human BMI was more correlated with LMA115 and BFT115 than other traits (e.g., DAY115, BW, LMD100) in pigs. This is in line with that BFT115 and LMA115 are in high genetic correlation with lean meat percentage (LMP) in pigs (51). We found that cattle body conformation traits had fewer overlapped genes with human BMI compared to pigs (Supplementary Figure 7, Figure 5A), revealing that pigs might be more desirable models for human body conformation traits than cattle.

**Figure 5.**
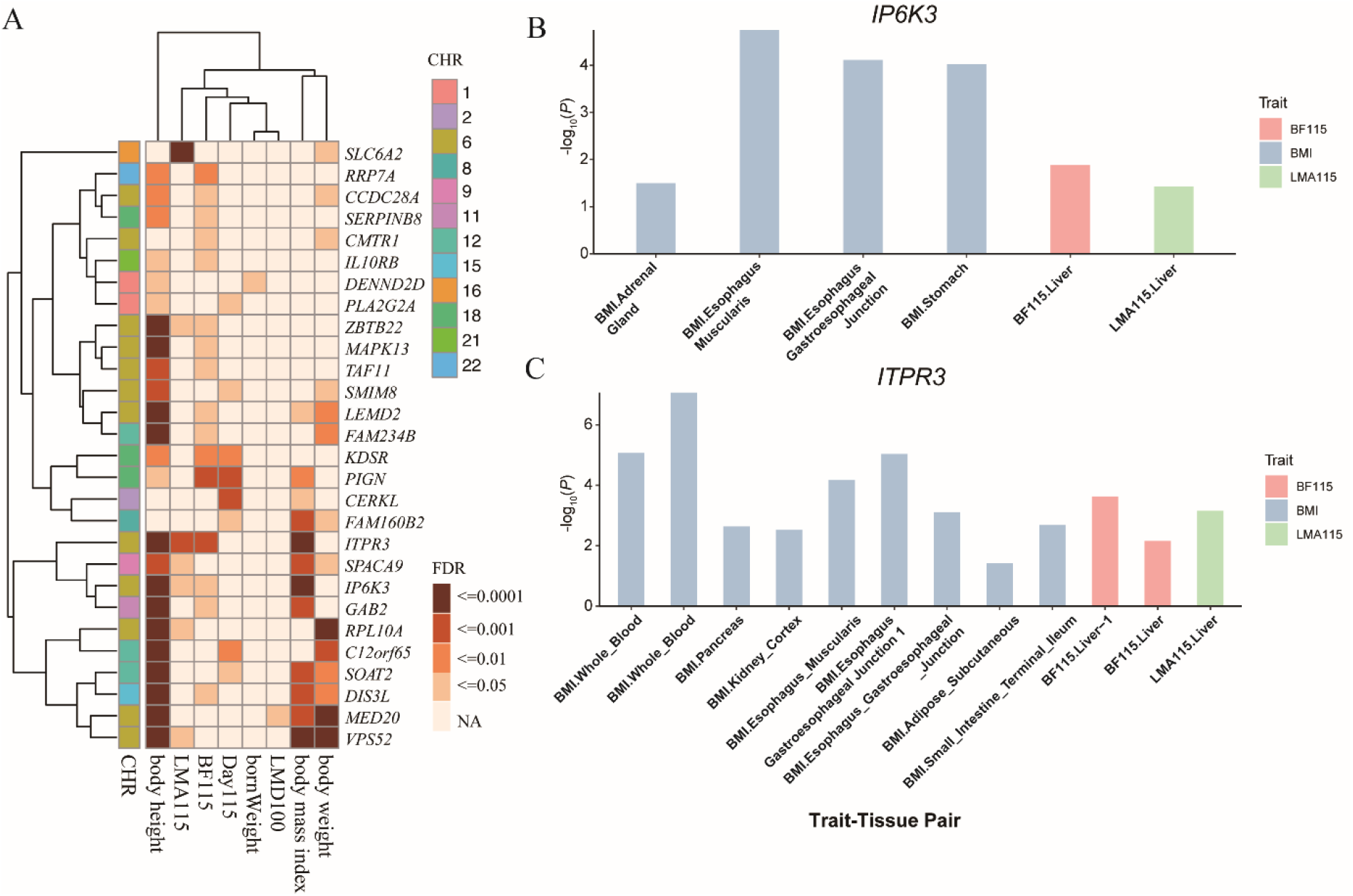
Results of *Case study 2*. A. The heatmap plot showed the lowest FDR in TWAS analysis of humans (BW, BH, BMI) and pigs (LMA115, BFT115, DAY115, bornWeight, LMD100) in the digestive system (esophagus muscularis, esophagus mucosa, esophagus gastroesophageal junction, colon transverse, colon sigmoid, small intestine terminal ileum, and liver of human. And small intestine, large intestine, duodenum, colon, ileum, jejunum, and liver of pig), the color of the box indicates the lowest FDR in corresponding tissues. B and C display the -log_10_(*P*-value) of *IP6K3* gene (B) and *ITPR3* gene (C) calculated from BMI, BFT115, and LMA115 across tissues. CHR: chromosome.

In the comparison of BFT115 and BMI, 747 out of 3,020 one-to-one orthologous genes being tested were significantly associated with BMI (Supplementary Figure 8B), and 23 were significantly associated with BFT115 (Supplementary Figure 8A). Six genes (i.e., *DIS3L, GAB2, IP6K3, ITPR3, PIGN, LEMD2*) were significantly associated with both BMI and BFT115. Interestingly, *ITPR3* and *IP6K3* were significantly associated with BFT115, LMA115, and BMI (Figure 5A-C, Supplementary Figure 8D, E). A previous study reported that the deletion of *IP6K3* protected mice from age-induced fat accumulation and insulin resistance (52). The methylation status of *ITPR3* might contribute to fat deposition (53). In addition, the animalQTLdb also reported that *ITPR3* was associated with body height, body weight, and body mass index significantly in pigs (54). Of note, even though the current sample size of BFT115 and LMA115 is limited (n _sample size_= 2,778), there are still significantly associated genes shared between human BMI and pig carcass traits, indicating that the genetics of similar phenotypes might be conserved across species. We believe that the FarmGTEx TWAS-server could help translate genetic results between breeds and species.

### Conclusions and future prospects

Here we presented the FarmGTEx TWAS-server to the research community. A unique feature of this TWAS-server is that it provides customized TWAS analysis and popular downstream functional annotation across multiple species, e.g., humans, cattle and pigs. The FarmGTEx TWAS-server can take individual genotype and GWAS summary statistics as input. As a result, it will output predicted gene expression and the TWAS results. It also supports to querying the existing TWAS results in the server by genes and traits. The case studies demonstrated that the FarmGTEx TWAS-server is effective for complex trait gene mapping and translating genetic findings across species.

Currently, there are three species (i.e., cattle, pigs, and humans) and their respective gene expression prediction models across a wide range of tissues implemented in the FarmGTEx TWAS-server. As the FarmGTEx project is developing, one promising direction of the FarmGTEx TWAS-server is to incorporate more tissues/cell types, molecular phenotypes (e.g., alternative splicing and enhancer expression), and species. We believe that the FarmGTEx TWAS-server will be a valuable resource that will help the entire community explore the genetic mechanism of complex traits and translate genetic results between breeds and species.

## DATA AVAILABILITY

The FarmGTEx TWAS-server is publicly available at http://twas.farmgtex.org. The genotype and gene expression data of animals is available at https://www.farmgtex.org/. The data for humans is available at https://www.gtexportal.org/. The code is available at https://github.com/ZhangZhenYang-zzy/TWAS.

## ACKNOWLEDGEMENTS

We thank all the researchers who have contributed to the publicly available data used in this research.

## Supplementary files

**Supplementary_Figure 1.**
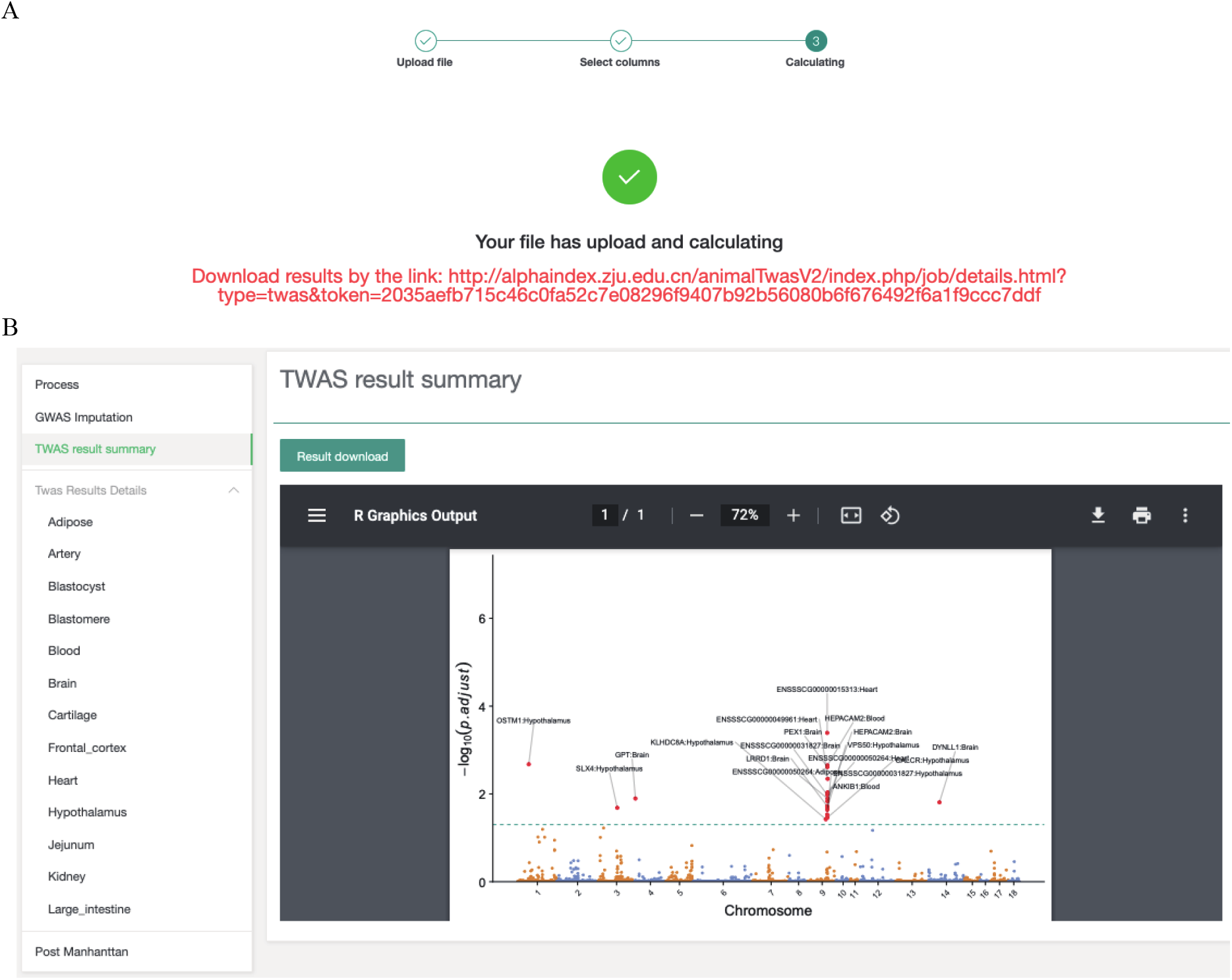
A. When users submit the job, the server will provide a link recording all the processes and results, which is in red font. B. The screenshot of the details of the successful job. The results include the GWAS imputation, TWAS results of each tissue. And the ‘Post Manhattan’ tab provides the interactive plot function.

**Supplementary_Figure 2.**
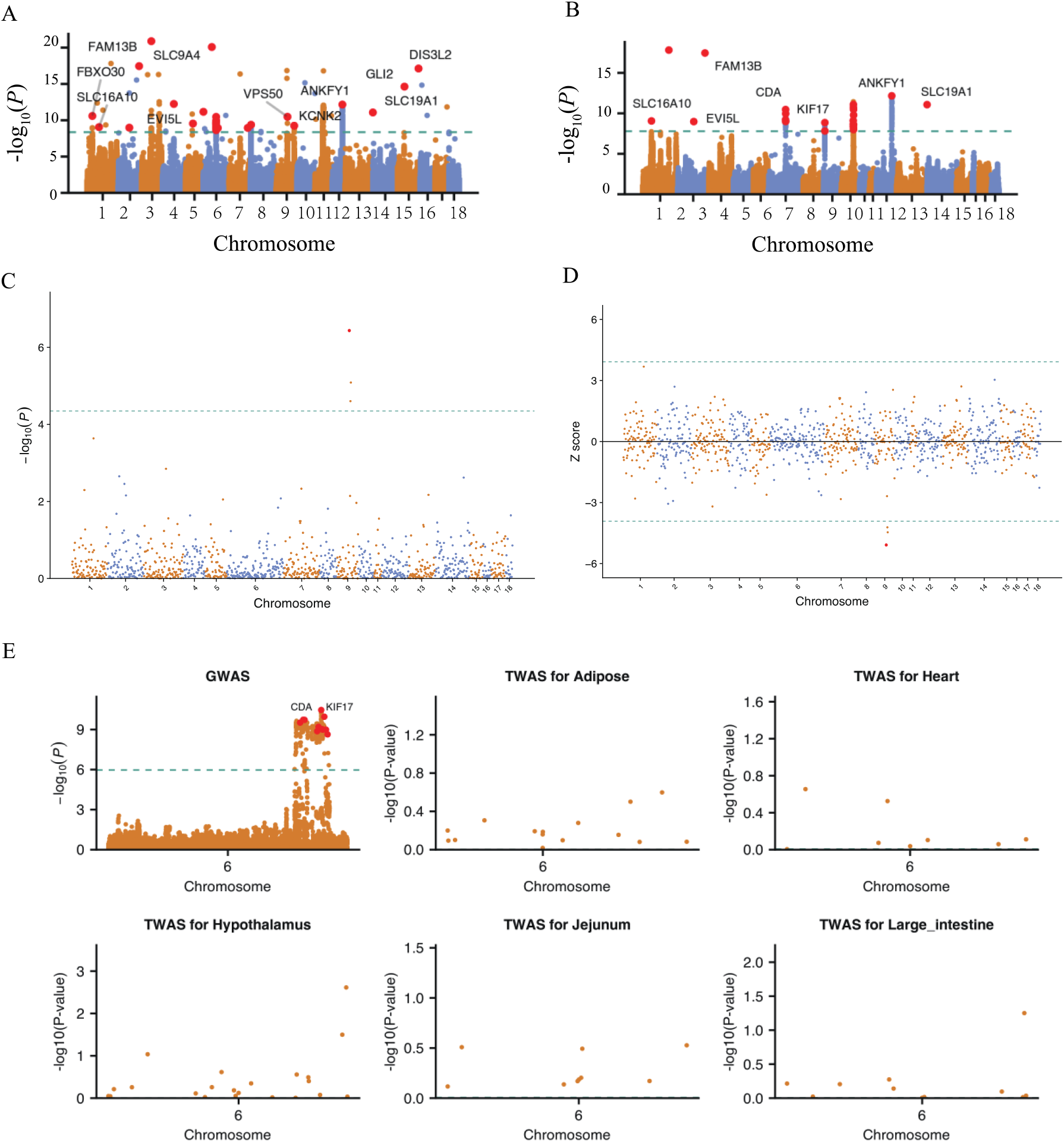
TWAS-server provides several kinds of Manhattan plots. A. Plots for GWAS. B. Plots for imputation GWAS. And plots for *P*-value(C) and z-score(D) of TWAS result in each tissue, (E) plots for a specific area by the “post-Manhattan” tab.

**Supplementary_Figure 3.**
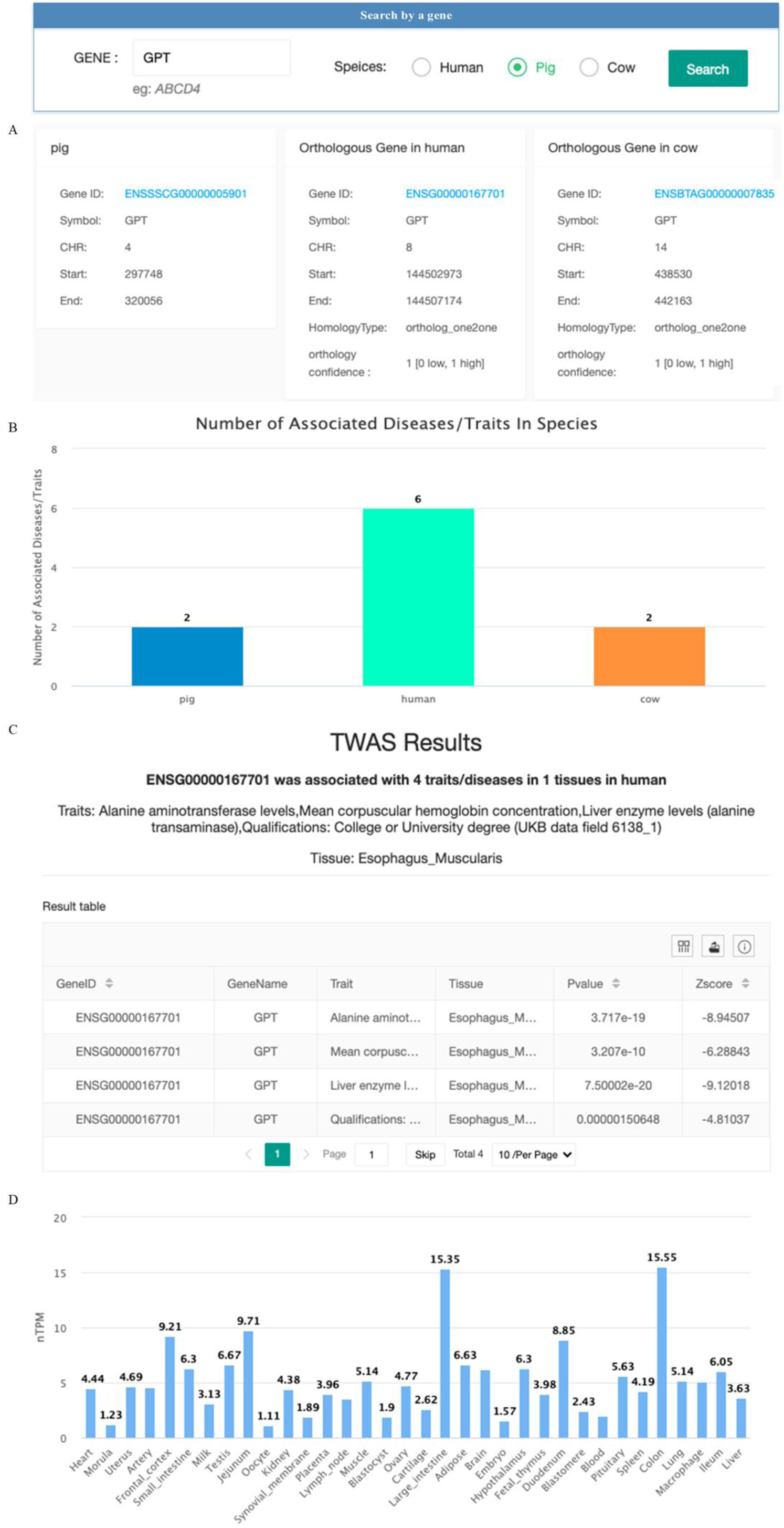
Operation flow for the ‘Search by gene’. A. The general information of the gene and orthologous gene in other species. B. The bar plot shows the number of associated trait-tissues in humans, pigs, and cattle. C. The details of TWAS summary statistics for interested species. D. And the expression level of the gene in humans, pigs, and cattle.

**Supplementary_Figure 4.**
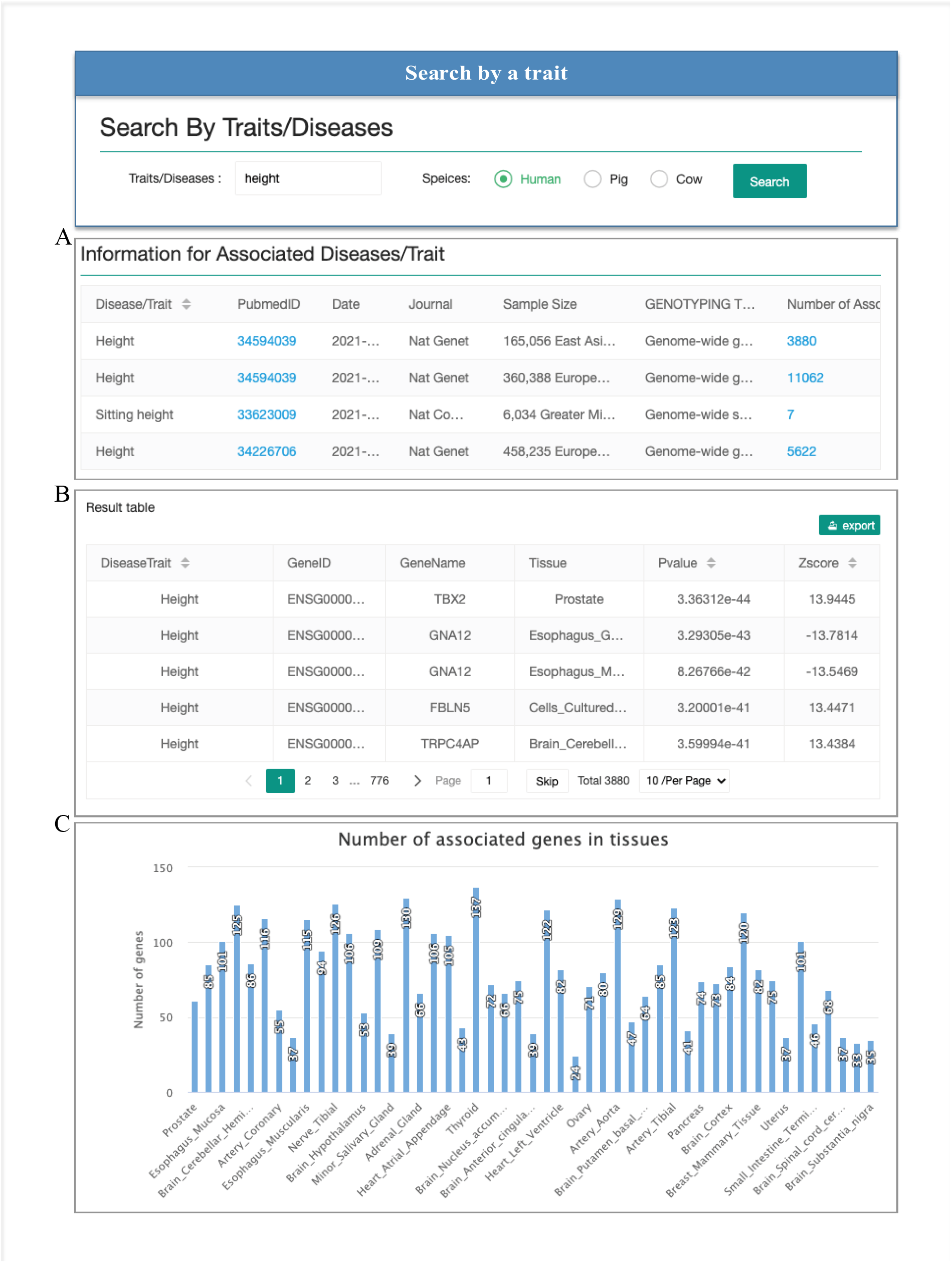
Screenshots of ‘Search by trait’. A. The number of the associated trait-tissues with the diseases/traits containing the searching keyword. B. By clicking the digital of the interesting study, a table displaying the TWAS summary statistics will be generated. C. A bar plot will show the number of associated genes across tissues.

**Supplementary_Figure 5.**
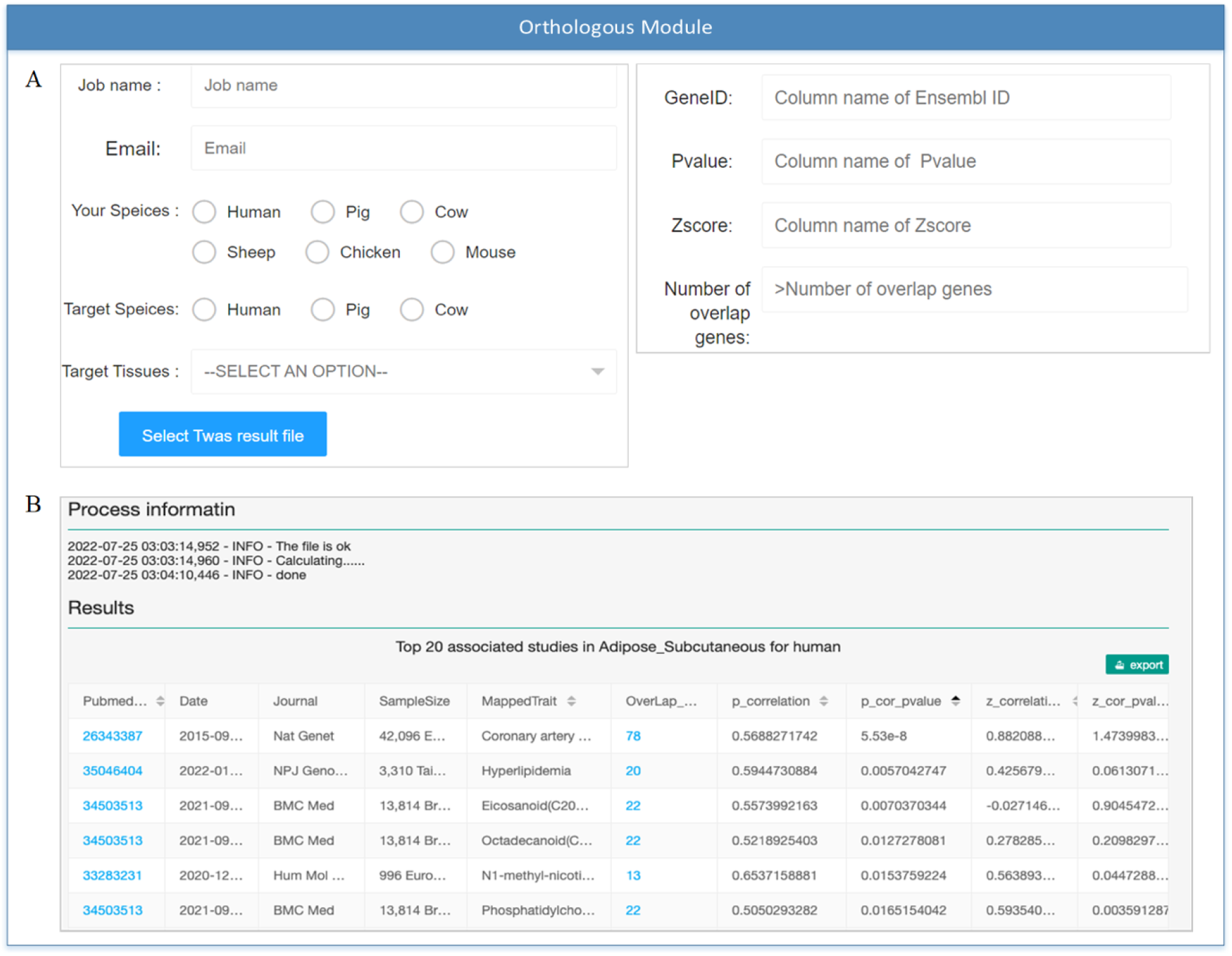
Screenshots of ‘Orthologous module’. A. Upload the GWAS summary statistics and select the options. B. The screenshot of the result.

**Supplementary_Figure 6.**
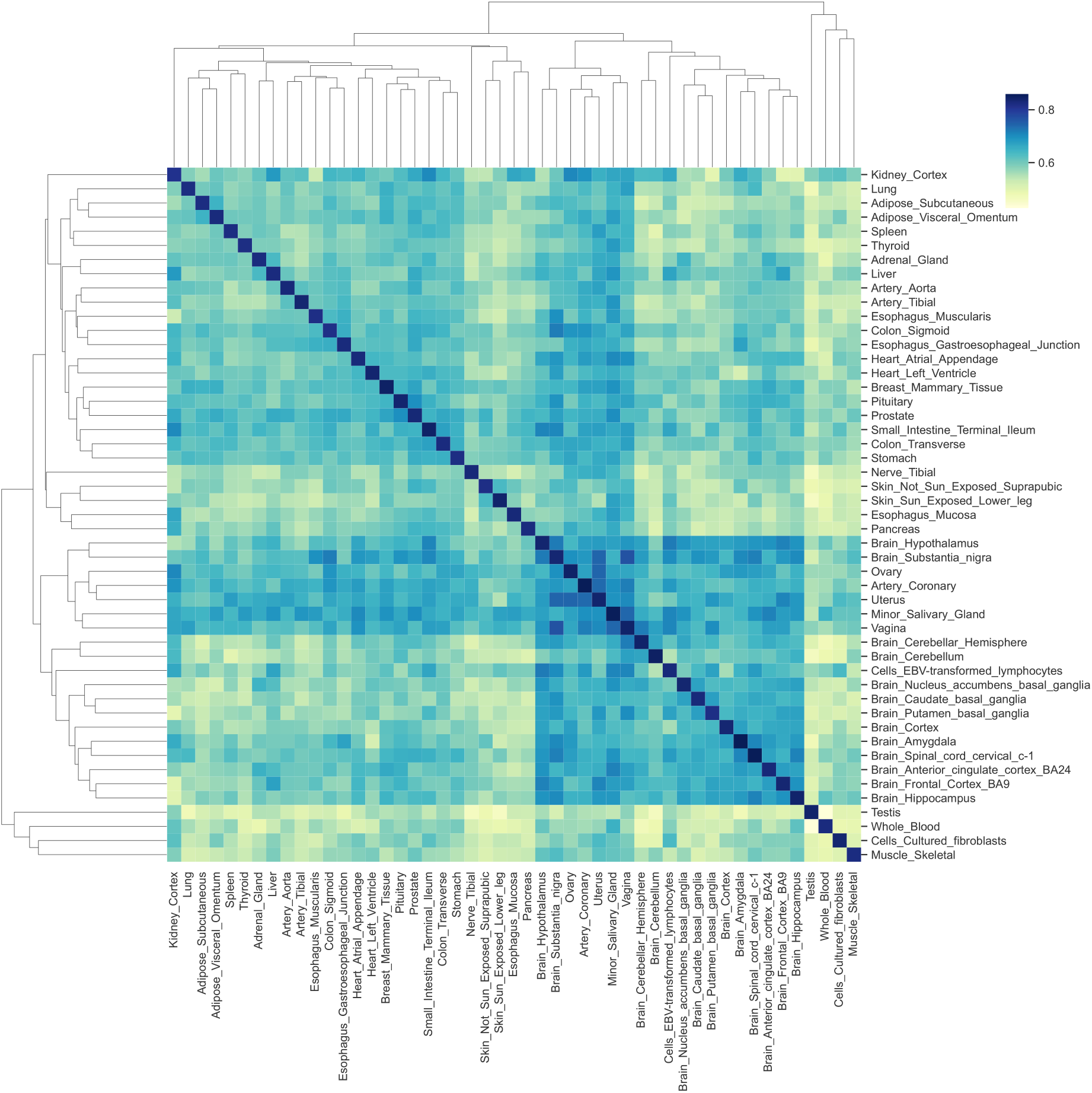
Cluster heatmap of correlation of the TWAS summary statistics across different tissues for BMI. The color of the box represents the Pearson correlation coefficient of *P*-value between two tissues.

**Supplementary_Figure 7.**
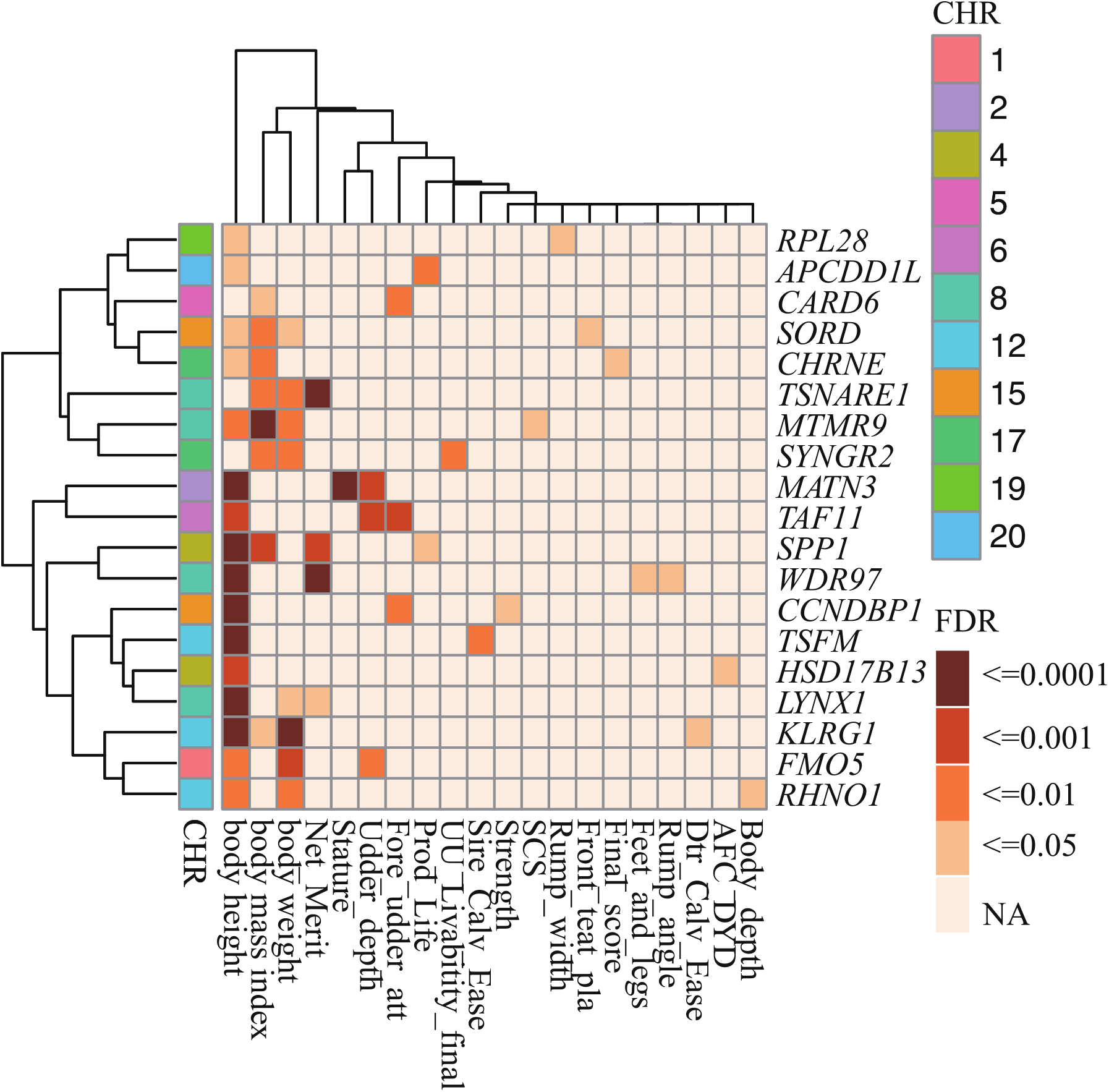
The heatmap showed the TWAS summary statistics of human (BW, BH, BMI) and cattle body conformation traits in the digestive system (esophagus muscularis, esophagus mucosa, esophagus gastroesophageal junction, colon transverse, colon sigmoid, small intestine terminal ileum, and liver of human. And rumen, jejunum, ileum, liver of cattle). The color means the *P*-value of the genes in different traits and tissues.

**Supplementary_Figure 8.**
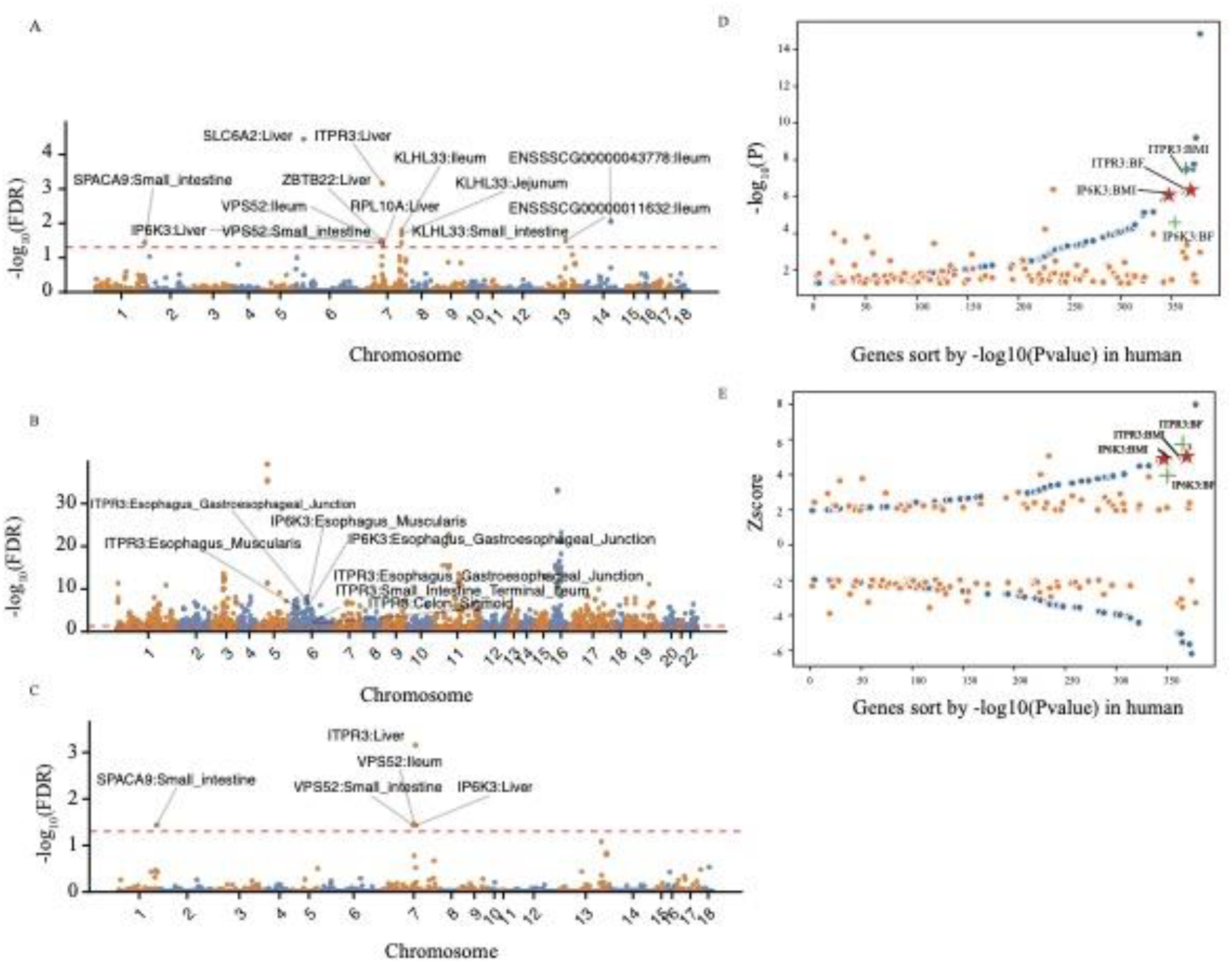
The significance of one-to-one orthologous genes between humans and pigs in TWAS analysis. A. The Manhattan plot of BMI TWAS results in humans. The Manhattan plot of BFT115 (B) and LMA115 (C) in the pig. D. On the x-axis, genes are ordered by the -log_10_(*P*-value) in human BMI, blue dots are the -log_10_(*P*-value) of human BMI, and the orange dots are the -log_10_(*P*-value) of pig BFT115. E. On the x-axis, genes are ordered by the -log_10_(*P*-value) in human BMI, and blue dots are the z-score of human BMI, and the orange dots are the z-score of pig BFT115.

**Supplementary Table 1**. The sample size of the RNA-seq samples for pig, and the number of the eVariants and eGENEs of the eQTL models. eGene: Genes with significant cis-eQTLs for each model. eVariants: Variants associated with at least one gene. etGene: tested genes for cis-eQTL. ePercent: Percentage of significant cis-eGenes in all tested genes.

**Supplementary Table 2**. The sample size of the RNA-seq samples for cattle, and the number of the eVariants and eGENEs of the eQTL models.

**Supplementary Table 3**. The sample size of the RNA-seq samples for human, and the number of the eVariants and eGENEs of the eQTL models.

**Supplementary Table 4**. The number of distinct eGenes detected by different methods in human, pig and cattle.

**Supplementary Table 5**. The number of distinct eVariants detected by different methods in human, pig and cattle.

**Supplementary Table 6**. The average heritability in each tissue and the average cross-validation R^2^ in prediction models for pigs.

**Supplementary Table 7**. The average heritability in each tissue and the average cross-validation R^2^ in prediction models for cattle.

**Supplementary Table 8**. The average heritability in each tissue and the average cross-validation R^2^ in prediction models for humans.

**Supplementary Table 9**. The tissue pairs used in the comparative analysis between humans and pigs. The “Number of orthologous genes” represents the eGenes shared in corresponding tissues. “Correlation of heritability” is the Pearson correlation of the orthologous genes’ heritability in corresponding tissues.

**Supplementary Table 10**. The tissue pairs used in the comparative analysis between humans and cattle. The “Number of orthologous genes” represents the eGenes shared in corresponding tissues. “Correlation of heritability” is the Pearson correlation of the orthologous genes’ heritability in corresponding tissues.

**Supplementary Table 11**. The tissue pairs used in the comparative analysis between pig and cattle. The “Number of orthologous genes” represents the eGenes shared in corresponding tissues. “Correlation of heritability” is the Pearson correlation of the orthologous genes’ heritability in corresponding tissues.

